# Mechanism of clinically relevant neurosecretory and metabolic impairment in Alzheimers and identification of engineered nanotherapy

**DOI:** 10.1101/2025.04.09.647859

**Authors:** Yutong Shang, Won Jin Cho, Seeya A. Munj, Siddhartha G. Jena, Suzan Arslanturk, Sushmita R. Patil, Xin Qi, Bhanu P. Jena

## Abstract

A central challenge in Alzheimer’s disease is understanding the mechanism of neuronal secretory dysfunction. By exploiting AI, we reveal how beta-amyloid disrupts key protein interactions within the neuronal secretory machinery. In a combinatorial strategy of reprogramming the neuronal secretory and metabolic components, we established a dual-target therapeutic framework that repairs synaptic and mitochondrial defects to counteract neurodegeneration in Alzheimer’s.

## Introduction

Alzheimer’s disease (AD) is characterized by progressive neurodegeneration and cognitive decline, driven by disruptions in neuronal secretion^1–3^ and mitochondrial homeostasis^4–6^. In the current study, using AlphaFold^7^ structural analysis, we reveal how beta-amyloid (Aβ) disrupts key protein interactions within the neuronal porosome, an essential secretory complex, compromising neurotransmitter release via the porosome^8–10^. Postmortem hippocampal analysis and AD neuronal cultures demonstrate a marked depletion of porosome proteins, including SNAP-25, Syntaxin-1A, and ATP1A3, establishing a mechanistic link between Aβ toxicity and synaptic failure. We previously showed that targeting oligomeric mitochondrial ATAD3A with the DA1 peptide^11,12^ rescued neuropathology and cognitive deficits in AD mice, restoring synaptic function and reducing mitochondrial oxidative stress. Notably, a combinatorial strategy integrating neuronal porosome reconstitution with DA1 achieves near-complete restoration of synaptic and metabolic homeostasis. Additionally, DA1 synergizes with the flavonoid apigenin to enhance neuronal viability and mitigate mitochondrial dysfunction. Our findings uncover an intrinsic connection between secretory and metabolic deficits in AD and establish a dual-target therapeutic framework that integrates synaptic repair and mitochondrial stabilization to counteract neurodegeneration.

Alzheimer’s disease (AD) is a devastating neurodegenerative disorder characterized by progressive cognitive decline, synaptic dysfunction, and widespread neuronal loss^13^. The pathogenesis of AD is closely linked to two fundamental disruptions in neuronal physiology: impaired neurotransmitter secretion and mitochondrial dysfunction. Synaptic failure, a hallmark of AD, results from deficits in the neuronal secretory machinery, leading to compromised neurotransmission and network connectivity. In parallel, mitochondrial dysfunction drives bioenergetic failure, oxidative stress, and neurodegeneration. While both pathways are implicated in AD pathology^14^, their interplay remains poorly understood, and effective therapeutic strategies that simultaneously target these dysfunctions are lacking.

The neuronal porosome is the universal secretory nanomachine responsible for neurotransmitter release, coordinating synaptic vesicle docking and fusion at the presynaptic membrane^8–10^. Recent evidence suggests that beta-amyloid (Aβ) disrupts key porosome components, including SNAP-25, Syntaxin-1A, and ATP1A3, leading to neurotransmission deficits. However, the precise molecular mechanisms underlying Aβ-induced porosome dysfunction remain elusive. Here, leveraging AlphaFold structural modeling and molecular analyses, we identify Aβ-induced structural perturbations within the neuronal porosome complex and demonstrate that reconstituting functional porosomes in AD neurons restores synaptic transmission.

Beyond synaptic deficits, mitochondrial dysfunction in AD is driven by pathological oligomerization of ATAD3A, a mitochondrial protein critical for fission-fusion balance and bioenergetics. ATAD3A oligomerization promotes mitochondrial fragmentation, disrupts ATP production, and exacerbates oxidative stress, ultimately contributing to neuronal death. We previously report that DA1, a peptide inhibitor of ATAD3A oligomerization^11,12^, restores mitochondrial function, enhances neuronal survival, and rescues cognitive deficits in AD mouse models^11,12^, supporting the notion that correcting mitochondrial damage could be a useful strategy to mitigate neurodegeneration.

Given the interconnected nature of secretory and metabolic dysfunctions in AD, we hypothesized that a combinatorial strategy targeting both pathways would provide superior therapeutic efficacy. Here, we demonstrate that porosome reconstitution, combined with DA1 peptide therapy, achieves near-complete restoration of synaptic and mitochondrial function in AD neurons. Furthermore, we identify the flavonoid apigenin as a metabolic enhancer that synergizes with DA1 to mitigate oxidative stress and improve neuronal viability. These findings establish a mechanistic framework linking synaptic and mitochondrial dysfunction in AD and provide a novel, dual-target therapeutic paradigm. By integrating structural biology, computational modeling, and functional assays, our study uncovers fundamental mechanisms underlying AD pathophysiology and advances a combinatorial therapeutic strategy with broad translational potential. This work highlights the importance of restoring neuronal secretory and metabolic networks as a viable path toward AD intervention.

## Result

### Reduction of neuronal porosome proteins in AD patient brains and AD cell culture

Western blot analysis of mouse neuroblastoma cells stably expressing either wild-type (APPwt) or Swedish mutant amyloid precursor protein (APPswe) demonstrate a reduction in key porosome proteins SNAP-25, Syntaxin-1A, and the sodium/potassium ATPase ATP1A3 (Fig. 1a). In postmortem hippocampal tissue from AD patients, immunofluorescence staining revealed a significant reduction in neuronal porosome proteins, SNAP-25, Syntaxin-1A, and ATP1A3, compared to normal subjects (Fig. 1b-e, S1a, S1b). These findings provided a basis for further investigation into whether SNAP-25, Syntaxin-1A, and ATP1A3 could serve as robust biomarkers for AD classification and, critically, how their interactions within the porosome complex might be disrupted in AD pathology.

**Figure 1.**
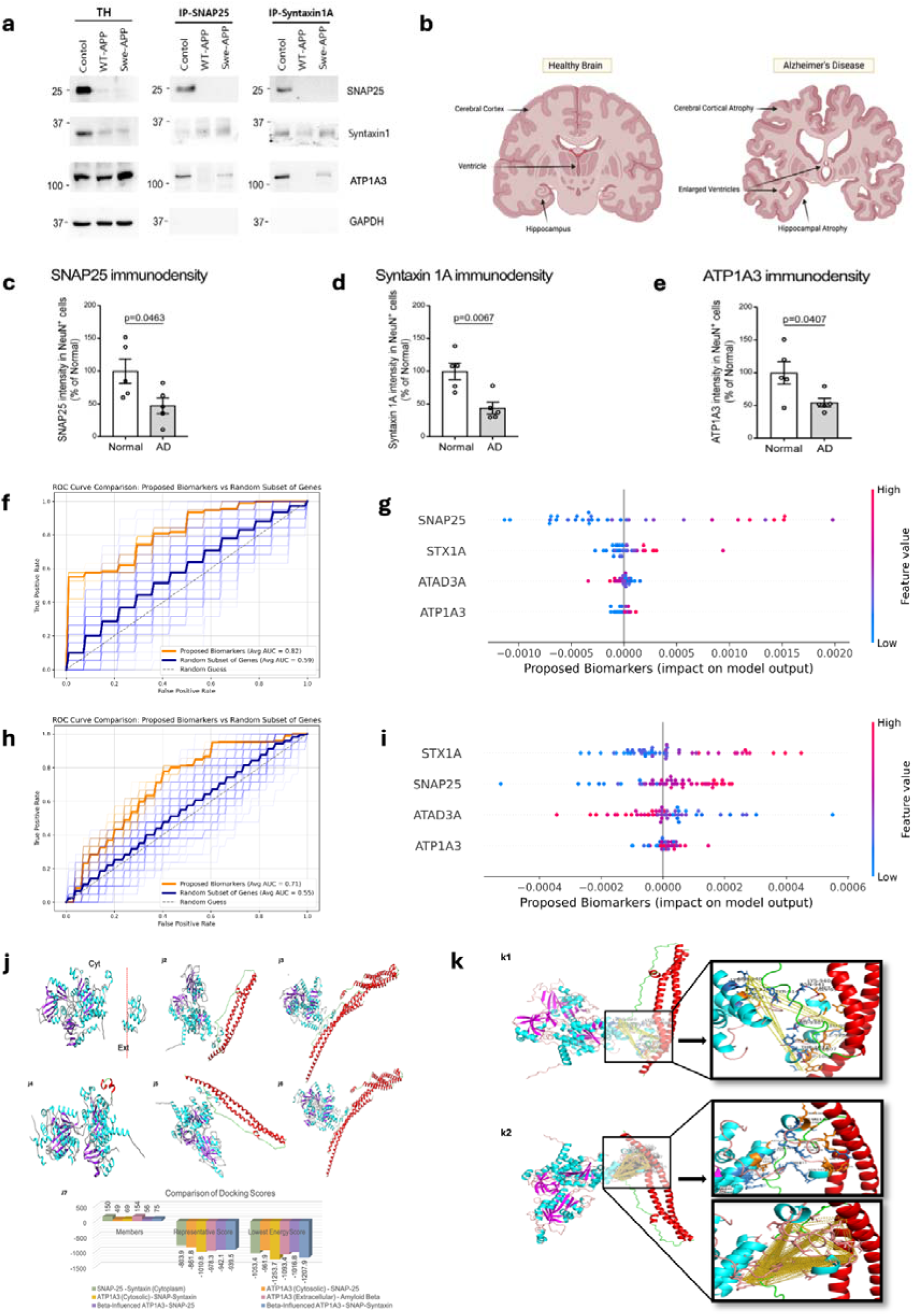
Immunocytochemistry and immunoblot demonstrate depletion of porosome proteins SNAP-25, Syntaxin-1A and ATP1A3 in AD mouse neurons and neurons in the hippocampus of AD patient postmortem brains. Studies using AI reveals altered binding interactions between porosome proteins SNAP-25, Syntaxin-1 and ATP1A3 when amyloid beta 1-42 peptide binding to the extracellular domain of ATP1A3. **(a)** Porosomes isolated from APP overexpressed AD neurons demonstrate depletion of porosome proteins SNAP-25, Syntaxin-1A and ATP1A3. Immunoblot demonstrate depletion of porosome proteins SNAP-25, Syntaxin-1A and Na+/K+-ATPase ATP1A3 in mouse neuroblastoma cells stably expressing either wild-type or Swedish mutant amyloid precursor protein APP. Notice the nearly undetectable levels of SNAP-25 protein in the immunoisolated porosome complex. **(b-e)** Postmortem hippocampus sections from control subjects and AD patients (n = 5 individuals/group) were stained with antibodies of (1) anti-SNAP25, Neurofilament heavy chain (NF-H), and NeuN; (2) anti-Syntaxin 1A, Synaptophysin, and NeuN; (3) anti-ATP1A3, MAP2, and NeuN. The intensity of SNAP25, Syntaxin 1A, and ATP1A3 in NeuN+ cells was quantified and shown in bar graphs, respectively. Hoechst was used to label nuclei. All data are presented as mean ± SEM. Statistical significance was determined by unpaired Student’s t-test. **(f-i)** Porosome proteins SNAP-25, Syntaxin-1A and ATP1A3, and the ATAD3A protein essential for mitochondrial fission and bioenergetics, provide superior classification performance in diagnosing AD over other gene products. **(f)** Receiver Operating Characteristic (ROC) curves for GSE5281, comparing model performance using selected biomarkers (ATP1A3, ATAD3A, SNAP25, STX1A) against randomly selected gene subsets. The model was run 100 times, selecting a random subset of genes in each iteration. The model trained on GSE5281 achieved an AUC of 0.82, significantly outperforming random gene subsets (AUC = 0.59), demonstrating the improved predictive capacity of the identified biomarkers. **(g)** SHAP-based biomarker importance for GSE5281. SHAP analysis confirms SNAP25 as the top predictor, with other biomarkers contributing to classification. Each gene’s SHAP score is derived from a normalized calculation where the total contribution across all genes sums to 1, providing insight into their relative influence on classification. The color gradient represents feature value distribution, with blue indicating lower expression levels and pink indicating higher expression levels, highlighting the impact of expression variability on model predictions. **(h)** ROC curves for GSE48350, evaluating classification performance on this dataset. The model was run 100 times, selecting a random subset of genes in each iteration. The targeted biomarkers (ATP1A3, ATAD3A, SNAP25, STX1A) yielded an AUC of 0.71, outperforming randomly selected gene subsets (AUC = 0.55), supporting the validity of these markers. **(i)** SHAP-based biomarker importance for GSE48350. The color gradient follows the same representation as in (b), indicating the relationship between expression levels and classification outcomes. **(j, k)** AI reveals altered binding interactions between porosome proteins SNAP-25, Syntaxin-1 and ATP1A3 when amyloid beta 1-42 peptide binding to the extracellular domain of ATP1A3. This binding of the amyloid beta peptide negatively impacts the binding interactions between ATP1A3-SNAP-25-Syntaxin-1A in the cytosolic domain within the neuronal porosome complex. **(j1)** Structure of ATP1A3, with cytosolic (Cyt.) and extracellular (Ext.) regions separated by a dashed line to indicate their distinct functional compartments. The extracellular region of ATP1A3 is exposed to the extracellular environment, where it interacts with amyloid beta, while the cytosolic region is involved in intracellular interactions with porosome proteins SNAP-25 and Syntaxin-1. **(j2)** The cytosolic domain of ATP1A3 (cyan and purple) favorably binds SNAP-25 (red and green). **(j3)** Interaction between the cytosolic domain of ATP1A3 (cyan and purple) and the SNAP-25+Syntaxin-1A complex (red and green), also favorably interact within the porosome machinery. **(j4)** Binding of amyloid beta (red and green) to the extracellular region of ATP1A3 (cyan and purple). Amyloid beta binds the extracellular region due to its location outside the cell, where it has been implicated in reducing the ATPase activity of ATP1A3 and neurodegeneration. The image includes the entire ATP1A3 protein, with the cytosolic region added for structural context. **(j5)** Binding of amyloid beta to the cytosolic domain of ATP1A3 (cyan and purple) and SNAP-25 (red and green), illustrating potential conformational changes in ATP1A3 induced by amyloid beta binding to its extracellular region. **(j6) The b**inding of amyloid beta to the cytosolic domain of ATP1A3 (cyan and purple) alters its binding with the SNAP-25+Syntaxin-1A complex (red and green) within the porosome**. (j7)** Comparison of docking scores for various interactions involving ATP1A3, SNAP-25, Syntaxin-1A, and amyloid beta (1-42) peptide. Results show that binding of amyloid beta (1-42) peptide to the extracellular domain of ATP1A3, results in tighter binding (lower energy scores) with SNAP-25 and reduced binding (higher energy scores) with the SNAP-25-Stntaxin-1 complex. This suggests that amyloid beta binding to the extracellular region of ATP1A3 may induce structural changes that enhance intracellular interactions with SNAP-25. These results suggest that while amyloid-beta binding enhances ATP1A3-SNAP-25 interactions, it negatively impacts the formation or stability of the larger ATP1A3-SNAP-25-Syntaxin-1A complex within the neuronal porosome, resulting in impaired secretion of neurotransmitters. **(k)** High resolution images of the detailed molecular interaction between ATP1A3 and SNAP25, with and without Amyloid-β binding. This figure illustrates the structural integration of ATP1A3 and SNAP25, highlighting residues within 5Å of the ligand. **(k1)** ATP1A3-SNAP25 Interaction: The interaction without Amyloid-β influence involves 19 key residues (*LEU 336, LYS 339, ARG 340, GLU 753, GLU 754, TYR 814, ARG 930, ARG 931, GLY 938, MET 939, LYS 940, ASN 941, LEU 997, ILE 998, GLY 1005, TRP 1006, VAL 1007, THR 1011, TYR 1012*). **(k2)** ATP1A3-SNAP25 Interaction in the presence of Amyloid-β: With Amyloid-β influence, the number of interacting residues increases to 28, including additional residues such as *CYS 333, LEU 757, ILE 758, ASN 761, LEU 762, GLU 815, THR 929, VAL 934, PHE 935, TYR 1013*. A notable difference in **(k2)** Amyloid-β-influenced ATP1A3-SNAP25 binding is the increase in hydrogen bonds (yellow dashed lines), which were identified based on a 1.8 Å distance threshold. This suggests stronger and potentially altered interactions in the presence of the Amyloid-β peptide. Hydrophilic (sky blue) and hydrophobic (orange) residues are also highlighted, to provide a clearer view, a zoomed-in version of the key residues and hydrogen bonding interactions.

### AI-powered insights into secretory defects in AD

The molecular architecture of the cell’s secretory portal, the porosome^8–10^, along with recent advancements in artificial intelligence (AI) and the development of AlphaFold^7^, offers a powerful approach to deciphering the mechanisms underlying neurotransmitter release defects in AD. Understanding these mechanisms could provide a pathway toward potential therapeutic interventions. To assess whether three key porosome-associated proteins, SNAP-25, Syntaxin-1A, and ATP1A3, serve as robust biomarkers for AD diagnosis, we developed a supervised autoencoder-based neural network designed to capture complex gene expression patterns while reducing dimensionality. The model employs an encoder-decoder architecture, where the encoder extracts a biologically meaningful low-dimensional representation of gene expression profiles to predict AD disease status. Trained on GSE5281, the model achieved an AUC of 0.82, significantly outperforming random gene subsets (AUC = 0.59), demonstrating the enhanced predictive power of SNAP-25, Syntaxin-1A, and ATP1A3 as AD biomarkers (Fig. 1f-i, S2).

Since amyloid beta (Aβ) has been found to alter the connectivity of olfactory neurons in the absence of amyloid plaques^19^, and SNAP25 is a potential target for early stage AD^20^, the binding of Aβ_(1-42)_ to the extracellular domain of the ATP1A3 protein, and its impact on the cytosolic facing domain of ATP1A3 with SNAP-25, and Syntaxin-1A interactions within the neuronal porosome complex was assessed using AI. This was examined using a multi-step computational workflow combining structure prediction, molecular docking, and structural analysis. This study revealed that binding interactions between porosome proteins SNAP-25, Syntaxin-1 and ATP1A3 exist, which is impaired upon binding of Aβ _(1-42)_ to the extracellular domain of ATP1A3 (Fig 1j,1k, S3). Results show that binding of Aβ_(1-42)_ peptide to the extracellular domain of ATP1A3, results in tighter binding (lower energy scores) with SNAP-25 and reduced binding (higher energy scores) with the SNAP-25-Syntaxin-1 complex. These results suggest that while Aβ_(1-42)_ binding enhances ATP1A3-SNAP-25 interactions, it negatively impacts the stability of the larger ATP1A3-SNAP-25-Syntaxin-1A complex within the neuronal porosome, resulting in impaired secretion of neurotransmitters.

This altered binding negatively impacting the porosome may also result in the loss of these porosome proteins as shown in Fig. 1a, resulting in neurodegeneration and progression of AD. These results, also provide for the first time, a molecular understanding of how the Aβ_(1-42)_ peptide negatively impacts neurotransmitter release and neuronal function, providing valuable insights into the treatment of AD. Since multiple proteins within the neuronal porosome complex are negatively affected or lost, introducing fully assembled, functional porosomes into AD neurons may be a more effective therapeutic approach than compensating for individual porosome proteins through overexpression or reconstitution. Furthermore, the structure of the molecular interacting domain of ATP1A3 where the Aβ_(1-42)_ binds, also provides a window for the development of engineered peptide biologics, those that could appropriately fold and bind with greater affinity with the Aβ_(1-42)_ peptide and neutralize it.

### Secretory and metabolic correctors for AD therapy

Porosomes are secretory portals at the cell plasma membrane where secretory vesicles transiently dock and fuse to expel a precise amount of intra-vesicular contents from the cell during secretion^8,9^. In neurons, porosomes are 15nm cup-shaped lipoprotein structures at the presynaptic membrane, composed of nearly 30 proteins^10^. A number of porosome proteins have previously been implicated in neurotransmission and neurological disorders, attesting to the crosstalk between porosome proteins and their coordinated involvement in release of neurotransmitter at the synapse. In AD, levels of porosome proteins CNPase (2,3-cyclic nucleotide phosphodiesterase) and the heat shock protein 70 (HSP70) are found to increase, while the levels of dihydropyrimidinase-related protein-2 (DRP-2) decrease^10^. Decreased levels of CNPase have been observed in the frontal and temporal cortex of patients with AD^21^. Similarly, porosome proteins SNAP-25 and synaptophysin are significantly reduced in neurons of patients with Alzheimer’s disease^22–24^. Mice that are SNAP-25 (+/-) show disabled learning and memory, and exhibit epileptic like seizures^25^. These results further support the conclusion that, alteration of one porosome protein impacts others within the complex, resulting in impaired porosome-mediated neurosecretion. This finding is similar to our recent studies on human bronchial epithelial (HBE) cells which shows that the ΔF508 cystic fibrosis transmembrane conductance regulator (CFTR) mutation in HBE cells, affects nearly a dozen porosome proteins including CFTR within the porosome complex. Therefore, the reprogramming by introduction of the porosome secretory machinery into the cell plasma membrane of Cystic Fibrosis (CF) cells, was able to restore normal mucin secretion and rescue from CF^26^. We therefore hypothesized that reconstitution of the 15 nm normal neuronal porosome complex in AD neurons would overcome secretory defects in neurotransmitter release.

Similarly, it is widely accepted that the mitochondrial production of reactive oxygen species (ROS) contributes to the detrimental alterations in the etiology and/or progression of many pathological conditions, including brain neurodegeneration^27^. Studies report that flavonoids can protect cells from different insults that lead to mitochondria-mediated cell death, and epidemiological data further show that some of these compounds attenuate the progression of diseases associated with oxidative stress and mitochondrial dysfunction^28^. Flavonoids are low molecular weight phenolic compounds, displaying significant ROS scavenging capability, including other cellular antioxidant effects^29–31^, and are hence ideal for the protection of brain neurons from the buildup of free radicals generated by the mitochondria leading to mitochondrial fission in AD.

ATAD3A is an ATPase family AAA-domain containing protein 3A found at mitochondrial contact sites. It spans the inner and outer mitochondrial membranes, with its N-terminal domain in the cytosol and C-terminal domain in the matrix^32–35^. ATAD3A regulates mitochondrial fission-fusion dynamics. Although global ATAD3A knockout (KO) is embryonic lethal^36^, selective ATAD3A KO in mice causes mitochondrial fragmentation and mitochondrial bioenergetics failure, resulting in cell death and tissue damage^37,38^. ATAD3A mutation is also associated with axonal neuropathy and spastic paraplegia^39–40^. Therefore, the proper function of ATAD3A is crucial for mitochondrial activity, integrity and cellular survival. In AD, ATAD3A undergoes oligomerization, resulting in Drp1-mediated mitochondrial fragmentation, leading to neurodegeneration^11^. Heterozygous knockdown of ATAD3A in AD transgenic mice abolishes ATAD3A oligomerization, reduces amyloid accumulation and neuroinflammation, and improves long-term memory^11^. These data further support the idea that aberrant ATAD3A oligomerization is a key pathogenic factor inducing AD neuropathology and further validates ATAD3A as a drug target.

We have developed DA1, a synthetic peptide that binds to ATAD3A and reduces ATAD3A oligomerization, which in turn decreases Drp1/ATAD3A binding that occurs during AD progression^11,12^. DA1 treatment reduces mitochondrial bioenergetic defects, and improves the survival rate of neurons exposed to toxic Aβ *in vitro*. DA1 penetrates the blood-brain-barrier, and is nontoxic with minimal effects on the immune response^11,12^. These physiological and pharmacodynamic properties make DA1 a promising candidate for AD therapy. We have tested^11^ the *in vivo* efficacy of DA1 in 5XFAD AD transgenic mice using a subcutaneous (SQ) Alzet mini-pump, delivering either the control peptide TAT or DA1 starting at 6 weeks of age show that 5XFAD mice treated with the control peptide exhibited enhanced ATAD3A oligomerization at 6 months of age, which was abolished by treatment with DA1^11^. Immunohistochemistry of 6-month-old 5XFAD mice brain revealed a significant increase in the density and area covered by Aβ. DA1 treatment significantly reduced this amyloid load^11^. Furthermore, DA1 decreased the immunoreactivity of GFAP (astrocytes) and Iba1 (microglia), suggesting reduced neuroinflammation. Additionally, DA1 treatment from 2 to 8 months of age improved the long-term memory of 5XFAD mice. DA1 treatment had no effect on the behavior of WT mice, further supporting the nontoxic nature of the peptide^11^.

In the current study in AD neuron culture, DA1 peptide demonstrated a partial rescue from AD both in cell viability and in reduced mitochondrial ROS production in neuronal cells exposed to Aβ (Fig. 2a-c, S5). However, the neuronal porosome reconstitution combined with the DA1 peptide of ATAD3A oligomers, was able to restore AD neurons viability to near normal levels and greatly reduced oxidative stress derived from mitochondria, demonstrating the potential for therapeutic applications. Additionally, the peptide and Apigenin, a small molecule flavonoid considered safe by the Food and Drug Administration (FDA) and used as a food additive, were also able to greatly increase viability in AD neurons and reduce to near normal levels mitoSOX (Fig. 2d-I, S6), also demonstrating its use in AD therapy. Collectively, these results demonstrate great promise in the combinatorial use of the porosome reconstitution secretory reprogramming and the DA1 peptide and Apigenin metabolic correctors, as highly safe and effective AD therapies. In agreement, our immunoblot analysis and immunocytochemistry study of human brain tissue (Fig. 1b), and the mouse neuroblastoma Neuro2a control cells, and our APPwt and APPswe AD cells (Fig. 1a), demonstrate significant decrease in SNAP-25 immunoreactivity and altered presence of other porosome proteins in both APPwt and APPswe neurons, confirming earlier reported studies of a decrease in SNAP-25 in patients with Alzheimer’s^23–25^. Consequently, in AD patients there is an increase in levels of SNAP-25 in the cerebrospinal fluid, which has been associated with cognitive decline^41,42^. Upon porosome reconstitution in Aβ treated neurons, our immunocytochemistry demonstrates the restoration of SNAP-25 and other porosome-associated proteins that result in increased cell viability to near normal levels and a decrease in mitoSOX intensity. In summary, we have tested the combinatorial use of the porosome, the small flavonoid molecule Apigenin, and our DA1 peptide, in treating AD. Results from our study demonstrate great promise in the combinatorial use of secretory (porosome) and metabolic (flavonoid and DA1 peptide) correctors for AD therapy, restoring AD neurons to near normal levels of viability and greatly reducing oxidative stress in the mitochondria.

**Figure 2.**
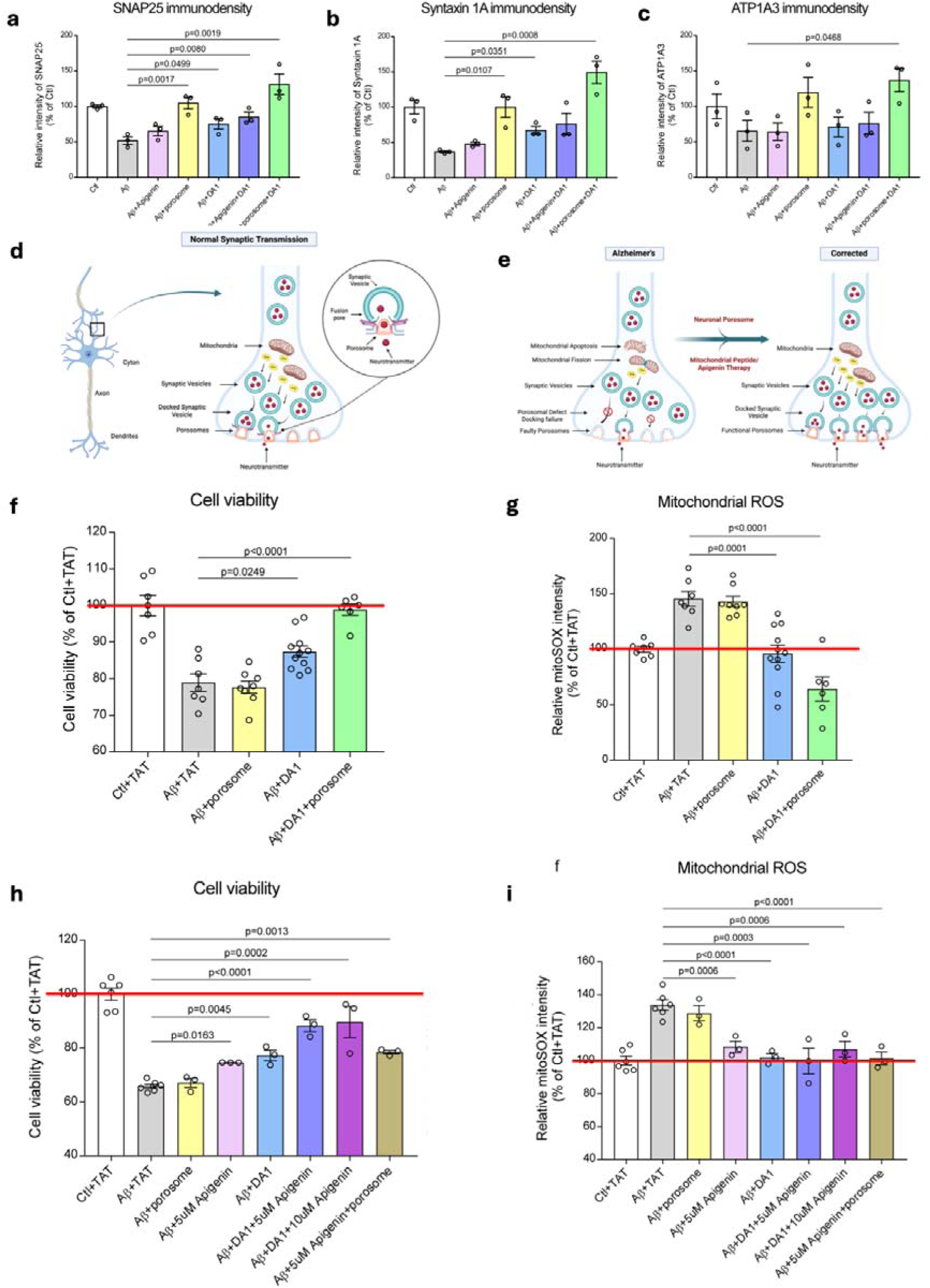
Immunocytochemistry demonstrating depletion of porosome proteins SNAP-25, Syntaxin-1A, and Na^+^/K^+^ Transporting ATPase alpha 3 (ATP1A3) in Alzheimer’s neurons (Aβ treated), which is fully restored to normal levels **(a-c)** following porosome-reconstitution therapy, and partially by Apigenin or the DA1 peptide. **(a-c)** The immuno densities of SNAP-25 **(a)**, Syntaxin-1A **(b)**, and ATP1A3 **(c)** in different treatment groups were quantified and shown in these bar graphs. At least 130 cells per group were analyzed, and the data were from 3 independent experiments. Data are presented as mean ± SEM. Statistical significance was determined by one-way ANOVA with Tukey’s post hoc test. **(d, e)** Schematic illustration of a synapse at the nerve ending in AD neurons, demonstrating abnormal mitochondrial function resulting in mitochondrial fission, loss in ATP generation, and altered porosome-mediated neurosecretion. **(f-i)** All of these defects in AD are seen to be overcome using porosome reconstitution and the DA1 peptide inhibitor of mitochondrial fission therapy. The combinatorial therapy of porosome reconstitution, the secretory corrector, and the DA1 peptide inhibitor of mitochondrial fission, the metabolic corrector, restores cell viability and reduces oxidative stress in mitochondria to normal levels in AD neurons. Similarly, combination therapy of the small flavonoid molecule Apigenin and the mitochondrial fission inhibiting linear DA1 peptide results in increase cell viability and reduction of mitochondrial oxidative stress to near normal levels in AD neurons.**(f)** Cell viability is significantly enhanced in Alzheimer’s neurons (Aβ+TAT) by porosome reconstitution in the presence of the DA1 peptide. Mouse hippocampal HT-22 neuronal cells were pretreated with porosome (175ng per well in a 96-well plate) overnight, followed by treatment with Aβ (oligomeric Aβ_1–42_ peptides) (5μM) together with DA1 (1μM) for 40h. Cell viability was measured by MTT assay after 16h of serum starvation. Note that the cell viability in Alzheimer’s neurons following porosome reconstitution in the presence of the DA1 peptide, demonstrate recovery to healthy neuron levels. Data represent the mean ± SEM from at least six independent experiments. Statistical significance was determined by one-way ANOVA with Tukey’s post hoc test. **(g)** Porosome reconstitution significantly reduces mitochondrial reactive oxygen species (ROS) production in Alzheimer’s neurons (Aβ+TAT) in the presence of DA1 peptide. HT-22 cells were pretreated with porosome (175 ng/well in a 96-well plate) overnight. Cells were then pre-incubated with DA1 (1 μM) for 1 hour, followed by treatment with oligomeric Aβ_1–42_ peptides (10 μM) for 12 hours. Mitochondrial ROS production was assessed 1 hour after the second peptide treatment using MitoSOX staining. MitoSOX intensity in Alzheimer’s neurons following porosome reconstitution in the presence of DA1 peptide demonstrates recovery to much reduced levels than found in healthy neurons. **(h)** Viability of HT-22 cells exposed to 5μM toxic oligomeric Aβ_1–42_ peptides mimicking Alzheimer’s (Aβ+TAT), is significantly enhanced when treated with 1 μM DA1 peptide and 5μM or 10μM Apigenin. Cells were treated with DA1 (1μM) + Aβ (5μM) + Apigenin (5uM or 10μM). After 24h, cells were washed using serum-free media, and again treated with DA1 (1μM) + Aβ (5μM) + Apigenin (5uM or 10μM) in serum-free medium. Cell viability was measured by MTT assay at 40h (16h serum starved). Data represent the mean ± SEM from at least three independent experiments. (**i)** Mitochondrial ROS production in Alzheimer’s neurons (Aβ+TAT) is significantly reduced in the presence of the DA1 peptide. However, not much change is observed in the presence of either 5μM or 10μM of Apigenin. HT-22 cells treated with DA1 (1μM) + Apigenin (5μM or 10μM) for 1h, and then exposed to Aβ (10μM) for 12h. Cells were treated again with DA1 (1uM) + Apigenin (5μM or 10μM) 1h before mitoSOX staining. Data represent the mean ± SEM from at least three independent experiments. Statistical significance was determined by one-way ANOVA with Tukey’s post hoc test.

## Discussion

This study lays the groundwork for therapies targeting both the secretory and metabolic dysfunctions in Alzheimer’s disease (AD)—a shift from conventional approaches focused primarily on amyloid clearance and inflammation. As of 2023, 187 clinical trials were evaluating 141 drugs for AD, with 36 in Phase 3, 87 in Phase 2, and 31 in Phase 1. Despite diverse mechanisms of action, most therapeutic strategies have centered on amyloid clearance, neurotransmitter modulation, or inflammation control^43^. In contrast, the current study takes a systems-level approach to correcting both the metabolic and secretory defects in AD at early stages of the disease, i.e., hyposmia or olfaction dysfunction. In >90% of AD patients, olfactory dysfunction precedes cognitive decline^19^, and the olfactory bulb of AD mice show a loss in expression of SNAP-25^20^. Since SNAP-25 levels in the cerebrospinal fluid (CSF) or in the blood of AD patients are significantly (p<0.0001) elevated, similar to the FDA-approved AD biomarker t-tau^45^, further supports SNAP-25 to serve as an AD biomarker. Dysfunction of odor discrimination can therefore also be used as a predictive measure for AD, and the proposed^45^ “Odorant Item Specific Olfactory Identification” test where certain odors can differentiate AD from general age-related decline in odor detection^46^ could also be used in combination with biomarkers and memory tests, to identify patients for the ‘Metabolic and Secretory Corrector AD Therapy.’

Delivery of the proposed therapy is also an important factor. Both the DA1 peptide and Apigenin can pass through the blood-brain barrier. However, since the olfactory bulb is connected via nerves to the brain centers involved in learning, memory and emotion, the DA1 peptide combined either with Apigenin or the porosome, deliverable through the nasal route, would be a viable and highly effective AD therapy, deliverable through the nasal route, especially at the early stages of the disease, would be a viable and highly effective AD therapy (Fig. 7). The therapy is scalable for commercialization since the DA1 peptide and Apigenin are readily available, and the scale-up of the neuronal porosome biologic could be achieved by immunoisolation from 3D cultures of the human neuroblastoma SH-SY5Y cell line grown in bioreactors^20^. 3D neuronal cultures will be used since the capacity of a neuron to establish synaptic connections is an essential prerequisite for its functional maturation, and that the formation of the synapse involves the appropriate assembly of the porosome secretory machinery at the nerve terminal (Fig. 2d). Since the neuronal porosome is present in all neurons, the above-mentioned scale-up approach offers reproducibility, reliability, and safety. One needs to be critically aware that, since three (Fig. 1) among possibly others, of the nearly 30 or so proteins of the neuronal porosome complex are impacted in Alzheimer’s, resulting in impairment function, one safe and highly effective approach would be to ameliorate the impairment by introducing normal neuronal porosomes at the pre-synaptic membrane. Porosome reconstitution therapy, therefore, remains the most optimal approach in ameliorating neurotransmitter release defects in AD. The normal secretion of neurotransmitter and its timely breakdown is physiologically relevant and optimal, as opposed to the available AD medications such as cholinesterase inhibitors that prevent the breakdown of acetylcholine in the brain to improve neuronal communication. Cholinesterase inhibitor drugs exhibit side effects that may include nausea, vomiting, diarrhea, muscle cramps, fatigue, and even weight loss^46^.

## MATERIALS AND METHODS

### Porosome Isolated and Reconstitution

*Neuro2a wt* cells were used to isolate porosomes for reconstitution. RIPA buffer (10 mM Tris-HCl, pH 8.0 • 1mM EDTA • 0.5mM EGTA • 1% Triton X-100 • 0.1% Sodium Deoxycholate • 0.1% SDS • 140mM NaCl • Dilute with dH_2_O) containing 0.1mM PMSF, 1mM ATP, and a cocktail of protease inhibitors, was used to lyse the cells. Protein in all fractions was estimated using BCA Protein assay Kit (ThermoFisher Cat. No. 23227, Rockford, IL 61101, USA). A total of 500 μg of the supernatant proteins were incubated with 2 μg of mouse monoclonal antibody raised against SNAP-25 (Santa Cruz, sc-390814) or Syntaxin1a (MilliporeSigma, SAB4502894) overnight, followed by incubation with 40 μl of 50% protein AG Magnetic agarose beads (Thermo Fisher., Cat. No. 78610) slurry for 1 h. The beads were washed with the binding/wash buffer (PBS with 150 mM NaCl) for 5 min at 4 °C with gentle agitation twice, then washed with deionized water once. The proteins were eluted with 100 uL of 0.1 M glycine, pH 2.2 and neutralized with 25uL of 1M phosphate buffer, pH 7.5. Porosome reconstitution into *Neuro2a APPwt and Neuro2a APPswe cells in culture* was achieved by exposing 0.125 μg/mL porosomes isolated from *Neuro2a wt* cells.

### Western Blot Analysis

Total cell homogenates (TH) and porosomes immuno-isolated (IP) from expanded WT (Control) mouse neuroblastoma cell culture and experimental neuroblastoma cells stably expressing either wild-type or Swedish mutant were subjected to SDS-PAGE and Western Blot analysis. 20μg of proteins (TH) and 20μl of isolated neuronal porosomes in Laemmli buffer were resolved in a 12.5% SDS-PAGE, followed by electrotransfer to 0.2-mm nitrocellulose (NC) membrane. The NC membrane was incubated for 1 hour at RT in blocking buffer (5% nonfat milk in PBS [pH 7.4] containing 0.1% Triton X-100 and 0.02% NaN_3_) and immunoblotted for 2 hours at RT with antibodies to SNAP-25 (1:1000, MilliporeSigma, S-9684) and GAPDH (1:3,000, MilliporeSigma, SAB600208). Anti-rabbit horseradish peroxidase secondary antibody conjugates were used (1:5,000, Cell Signaling Technology Cat. No. 7074), and then developed with SuperSignal™ West Femto Maximum Sensitivity Substrate (Thermo Fisher, Cat. No. 34095) and exposed using by ChemiDoc XRT+ image system (Bio-Rad, Richmond, CA 94806, USA).

### AI driven model development and biomarker analysis

We developed a supervised autoencoder-based neural network that effectively captures complex gene expression patterns while reducing dimensionality. The model consists of an encoder-decoder architecture, where the encoder learns a low-dimensional latent representation of gene expression profiles, preserving the most biologically relevant information. The decoder reconstructs the original input to ensure retention of key signal characteristics. The encoded representation is subsequently passed to a classification head that predicts disease status (AD or control). Training was performed using all genes in the dataset, allowing the network to learn a comprehensive feature set. Following initial model training, we employed SHapley Additive exPlanations (SHAP) (Lundberg, S. M., & Lee, S. I. (2017). A unified approach to interpreting model predictions. Advances in Neural Information Processing Systems, 30) to quantify the contribution of individual genes to classification performance. SHAP assigns an importance score to each feature, helping to identify the most influential biomarkers in distinguishing AD from control samples. The color gradients in the SHAP summary plots provide an additional layer of interpretability, illustrating how high and low expression levels of specific genes correlate with disease prediction. To evaluate the predictive value of the targeted biomarkers, we conducted a validation classification experiment where a separate model was trained exclusively on the selected genes. This model was benchmarked against 100 independent runs using randomly selected gene subsets of the same size. The goal was to determine whether the chosen biomarkers provided superior classification performance compared to arbitrary selections, thus validating their biological and diagnostic relevance.

### Dataset-Specific Results

The first dataset, GSE48350, is enriched for AD-related neuropathology and consists of 173 control and 80 disease samples. When trained on all available genes, the autoencoder model achieved 90.2% accuracy, confirming that gene expression patterns in this dataset provide strong discriminatory power. After applying SHAP to identify the influence of the targeted biomarkers (ATP1A3, ATAD3A, SNAP25, STX1A), a separate model was trained exclusively on the selected biomarkers. This model achieved an AUC of 0.71, outperforming randomly selected gene subsets, supporting the robustness of these features in disease classification. The second dataset, GSE5281, provides a broader sample distribution with 74 control and 87 disease samples. The autoencoder model achieved 87.9% accuracy when trained on the full transcriptome. SHAP analysis revealed SNAP25 as a highly influential gene. A separate classifier using only the targeted biomarkers achieved an AUC of 0.82, again surpassing models trained on randomly selected gene subsets. The SHAP-based interpretation confirms the biological relevance of these selected genes, particularly SNAP25, whose expression pattern strongly correlates with AD status. Taken together, these results demonstrate that data-driven feature selection via SHAP improves the interpretability of deep learning models applied to transcriptomic data. The ability of these biomarkers to generalize across different datasets highlights their potential utility in understanding AD pathology and developing predictive models for early detection.

### Protein-protein interaction approach

To investigate the impact of amyloid beta (1-42) binding on the ATP1A3-SNAP-25-Syntaxin-1A interactions within the neuronal porosome complex, a multi-step computational workflow combining structure prediction, molecular docking, and structural analysis was employed.

#### Protein Structure Prediction and Preparation

The structural models of ATP1A3, SNAP-25, and Syntaxin-1A were obtained using **AlphaFold-Multimer (v2.3.1)**, a state-of-the-art deep learning-based protein structure prediction tool that leverages multiple sequence alignments and structural templates to generate accurate multi-chain protein complexes^7^.ATP1A3 was modeled separately for its cytosolic and extracellular domains to evaluate their respective binding interactions. The predicted structures were visually inspected and processed using **PyMOL (v2.5)**^48^ ensuring correct domain orientation and structural integrity. Minor inconsistencies in loop positioning and atomic clashes were assessed visually, and the structures were prepared for downstream docking.

#### Molecular Docking Simulations

Protein-protein interactions were evaluated through rigid-body docking simulations using the **ClusPro** docking server, which applies a Fast Fourier Transform (FFT)-based sampling algorithm to predict energetically favorable binding conformations^49^. ATP1A3 was treated as the receptor in all docking experiments, while SNAP-25, Syntaxin-1A, and amyloid beta (1-42) peptides were treated as ligands in independent docking runs. The docking was conducted in two scenarios: (1) ATP1A3 cytosolic domain docking with SNAP-25 and SNAP-25-Syntaxin-1A complex, and (2) amyloid beta binding to the extracellular domain of ATP1A3, followed by intracellular docking analysis to assess structural perturbations. The docking poses were ranked based on ClusPro’s default scoring function, which combines van der Waals, electrostatics, and desolvation energies.

#### Binding Energy Analysis and Structural Interpretation

Docking scores from ClusPro were analyzed by comparing the weighted docking scores and lowest energy conformations across different interaction scenarios. The structural alignment and visualization of docking poses were performed using PyMOL (v2.5) to assess conformational changes and binding interface stability. Quantitative comparisons of docking scores were interpreted to infer the effect of amyloid beta on ATP1A3-SNAP-25 interactions.

### AD-associated cell Cultures

Mouse hippocampal HT-22 cells (MilliporeSigma, SCC129) and mouse neuroblastoma cell line Neuro2a cells (ATCC, CCL-131) were cultured in DMEM supplemented with 10% (v/v) heat-inactivated FBS and 1% (v/v) antibiotics (100 unit/mL penicillin, 100 μg/mL streptomycin). Neuro2a cells stably overexpressing human APP wildtype (APPwt) or Swedish mutant (APPswe, K670N and M671L APP, clone Swe.10) obtained from Dr. Gopal Thinakaran (University of Chicago), were cultured as previously described^11^. Cells were plated on poly-D-lysine/laminin (P6407, Sigma-Aldrich)-coated culture plates with or without coverslips at an appropriate cell density. All cells were maintained at 37 °C and 5% CO_2_.

### Control TAT and DA1 Peptides

Control peptide TAT and DA1 peptide (Product number P103882, Lot# 0P082714SF-01) were synthesized at Ontores (Hangzhou, China). Their purities were assessed as >90% by mass spectrometry. Lyophilized peptides were dissolved in sterile water and stored at −80 °C until use.

### MTT Assay

MTT (3-(4,5-dimethylthiazol-2-yl)-2,5-diphenyltetrazolium bromide) assay is a colorimetric method that measures cell viability, proliferation, and cytotoxicity. It is based on the reduction of a yellow tetrazolium dye, MTT, into purple formazan crystals by metabolically active cells. Cell viability was measured using an in vitro toxicology assay, MTT-based kit (Roche, 11465007001), according to the manufacturer’s instructions.

### MitoSOX staining

Cells cultured on 96-well plates were washed with PBS and then incubated with 5 μM MitoSOX™ Red (Invitrogen, M36008), a mitochondrial superoxide indicator, for 10 min at 37 °C. MitoSOX intensity was measured (510 nm excitation/580 nm emission) by infinite M1000 multimode fluorescence plate reader (Tecan, Switzerland). The MitoSOX fluorescence density was normalized to the fluorescence density of DAPI (355 nm excitation/460 nm emission) which stained the nuclei.

### Immunofluorescence Cytochemistry

Cells were grown on coverslips, fixed with 4% paraformaldehyde for 20 min at room temperature, permeabilized with 0.1% Triton X-100 in PBS, and blocked with 2% normal goat serum. The cells were incubated with the indicated primary antibodies overnight at 4 [. After washing with PBS, the cells were incubated with Alexa Fluor goat anti-mouse/rabbit 568/488 secondary antibody (1:1000; Thermo Fisher Scientific) for 2 h at room temperature. The nuclei were counterstained with Hoechst dye (1:10,000; Sigma-Aldrich). Coverslips were mounted and slides were imaged by a confocal microscope (Olympus, Tokyo, Japan; Fluoview FV3000).

For immunofluorescence staining of postmortem brains of normal subjects and AD patients, paraffin-embedded sections were permeabilized with 0.2% Triton X-100 in PBS buffer, followed by blocking with 5% normal goat serum. The brain sections were incubated with the indicated primary antibodies overnight at 4 °C and then stained with Alexa Fluor goat anti-mouse/rabbit/guinea pig 568/488/633 (1:500) for 2 h at room temperature. The nuclei were counterstained with Hoechst dye (1:10,000; Sigma-Aldrich). Images of the staining were acquired using a confocal microscope (Olympus).

All quantification of immunostaining was performed using ImageJ software. The same image exposure times and threshold settings were used for all sections from all the experimental groups. Quantitation was performed blinded to the experimental groups.

### Reagents

Apigenin (S2262) was purchased from Selleck Chemicals LLC (Houston, TX, USA).

### Preparation of oligomeric Aβ_1–42_

The Aβ_1–42_ (GenicBio Limited) peptides were dissolved in 1,1,1,3,3,3-hexafluoro-2-propanol (HFIP; 105228, Sigma-Aldrich) to a final concentration of 5 mM and placed in a chemical hood overnight. The next day, HFIP was further evaporated using a SpeedVac concentrator for 1 h. Monomer Aβ (5 mM) was prepared by dissolving Aβ peptide in anhydrous dimethyl sulfoxide (Sigma-Aldrich). The oligomeric Aβ peptides were prepared by diluting the monomer Aβ solution in Dulbecco’s Modified Eagle Medium (DMEM)/F12 and then incubating at 4 °C for 24 h.

### Human postmortem brain samples

All postmortem brain samples were collected by the National Institutes of Health (NIH) NeuroBioBank (NBB; https://neurobiobank.nih.gov/) under the approval of the Institutional Review Boards (IRB) and the institution’s Research Ethics Board. The human postmortem brain samples used in the experiments were obtained from the NBB under a material transfer agreement (MTA) between the NIH and Case Western Reserve University and listed in Supplementary Table 1.

### Content

A small part of this manuscript has previously been published as a preprint in *bioRxiv* ^50^.

## Contributions

The idea for the use of porosome-reconstitution therapy was developed at Porosome Therapeutics Inc. The research design was developed by B.P.J. S.G.J. serving as a consultant for Porosome Therapeutics, Inc., identified and assessed the requirement for the use of AI approaches to determine altered protein-protein interactions within the porosome complex, which helped elucidate the possible mechanism of secretory defects in AD. W-J.C isolated the porosomes and conducted Western blot analysis, identifying altered porosome in AD neurons. YTS conducted the immunocytochemistry and assessed cell viability and mitochondrial toxicity under the expert guidance of XQ at Case Western Reserve University as a paid contract from Porosome Therapeutics, Inc. S.A.M conducted AI studies and developed approaches under the expert guidance of S.A. at Wayne State University as a paid consultant from Porosome Therapeutics, Inc. SP created the schematics, assembled the figures, and formatted the manuscript. B.P.J. and SP wrote the paper. All authors participated in critical reading and discussion of the manuscript.

## Acknowledgments

The work presented in this article was supported by Porosome Therapeutics, Inc., and the Viron Molecular Medicine Institute, Boston, MA.

## SUPPLEMENTARY FIGURES

**Table I:** Antibodies.

| Antibody | Manufacture | Catalog # | Dilution |
| --- | --- | --- | --- |
| Synaptophysin | Abcam (Cambridge, UK) | ab32127 | 1.430555556 |
| Neurofilament (NF-H) | Sigma-Aldrich (St. Louis, MO, USA) | N4142 | 0.736111111 |
| β-actin | Sigma-Aldrich (St. Louis, MO, USA) | A1978 | 1:10000 |
| MAP2 | Cell Signaling (Danvers, MA, USA) | 4542 | 0.388888889 |
| NeuN | MilliporeSigma (Burlington, MA, USA) | ABN90 | 0.388888889 |
| SNAP25 | BioLegend (San Diego, CA, USA) | 805002 | 0.180555556 |
| Syntaxin 1A | BioLegend (San Diego, CA, USA) | 827002 | 0.388888889 |
| ATP1A3 | Novus Biologicals (Centennial, CO, USA) | NB300-540 | 0.180555556 |

**Table II:**
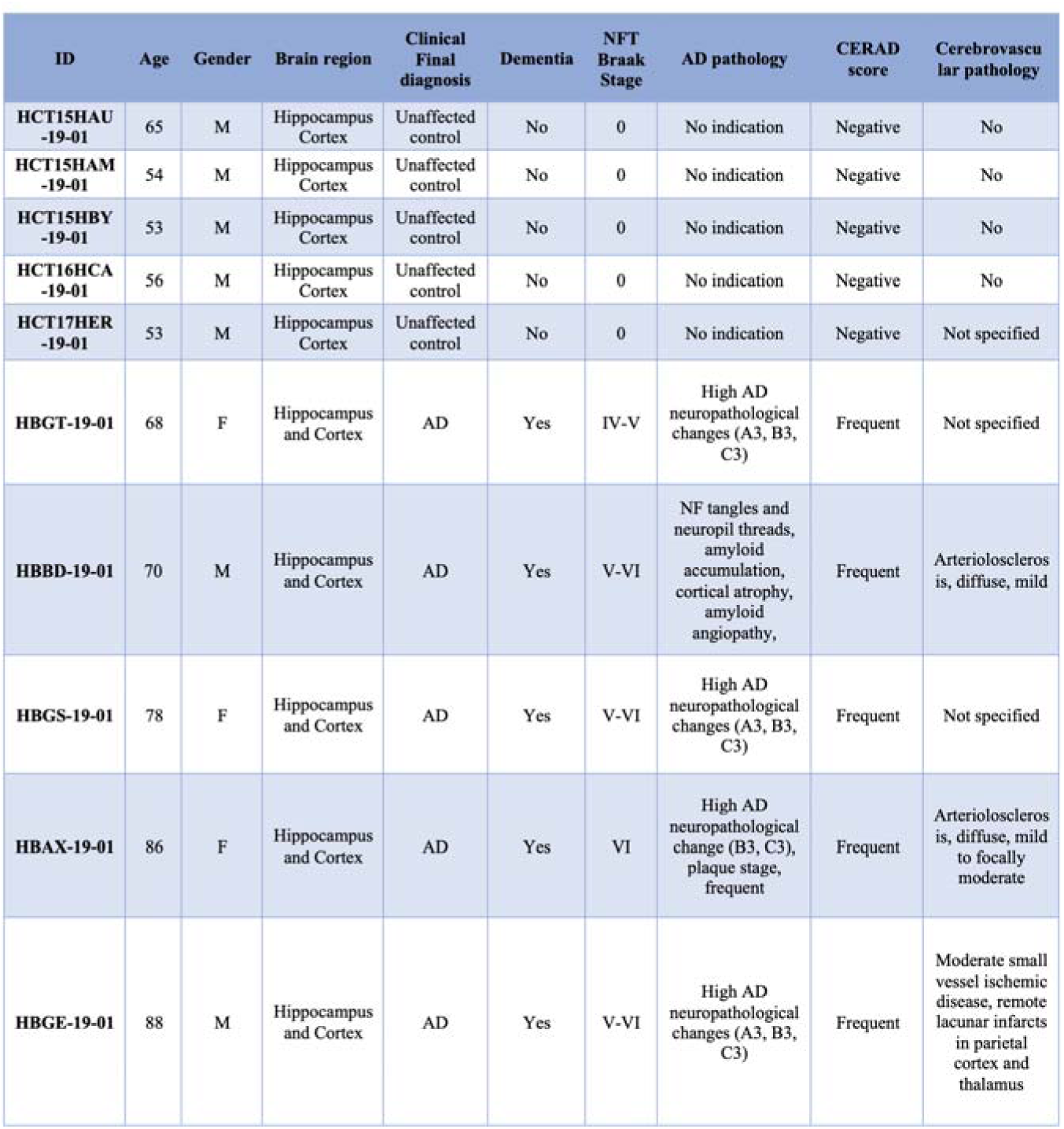
Human postmortem brain samples.

**Figure S1.**
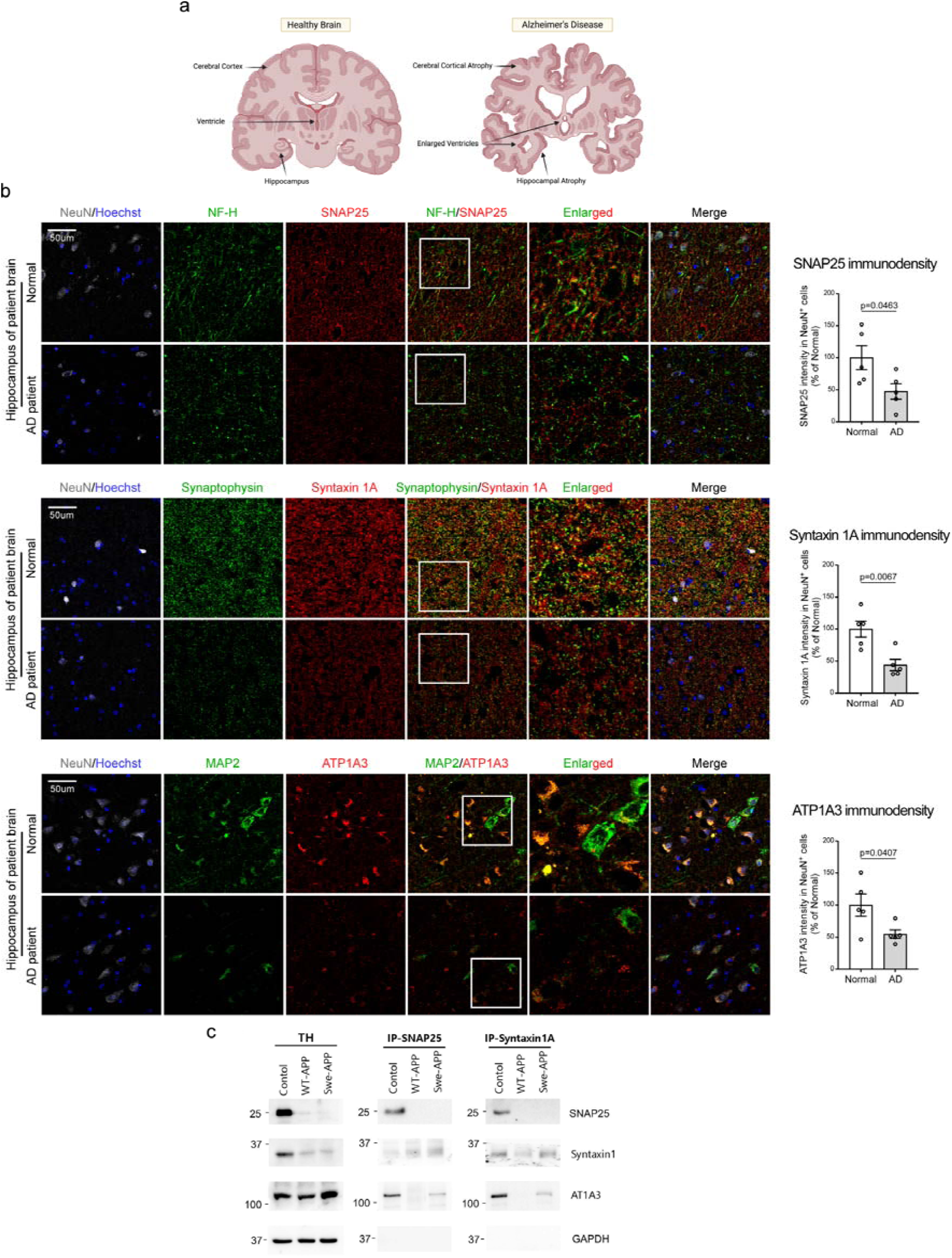
Immunocytochemistry and immunoblot demonstrate depletion of porosome proteins SNAP-25, Syntaxin-1A and ATP1A3 in the hippocampus **(a)** of AD patient postmortem brains **(b)**, and porosomes isolated from APP overexpressed AD neurons **(c)**. **(b)** Postmortem hippocampus sections from control subjects and AD patients (n = 5 individuals/group) were stained with antibodies of (1) anti-SNAP25, Neurofilament heavy chain (NF-H), and NeuN; (2) anti-Syntaxin 1A, Synaptophysin, and NeuN; (3) anti-ATP1A3, MAP2, and NeuN. The intensity of SNAP25, Syntaxin 1A, and ATP1A3 in NeuN+ cells was quantified and shown in bar graphs, respectively. Hoechst was used to label nuclei. All data are presented as mean ± SEM. Statistical significance was determined by unpaired Student’s t-test. **(c)** Immunoblot demonstrate depletion of porosome proteins SNAP-25, Syntaxin-1A and Na+/K+-ATPase ATP1A3 in mouse neuroblastoma cells stably expressing either wild-type or Swedish mutant amyloid precursor protein (APPNotice the nearly undetectable levels of SNAP-25 protein in the immunoisolated porosome complex.

**Figure S2.**
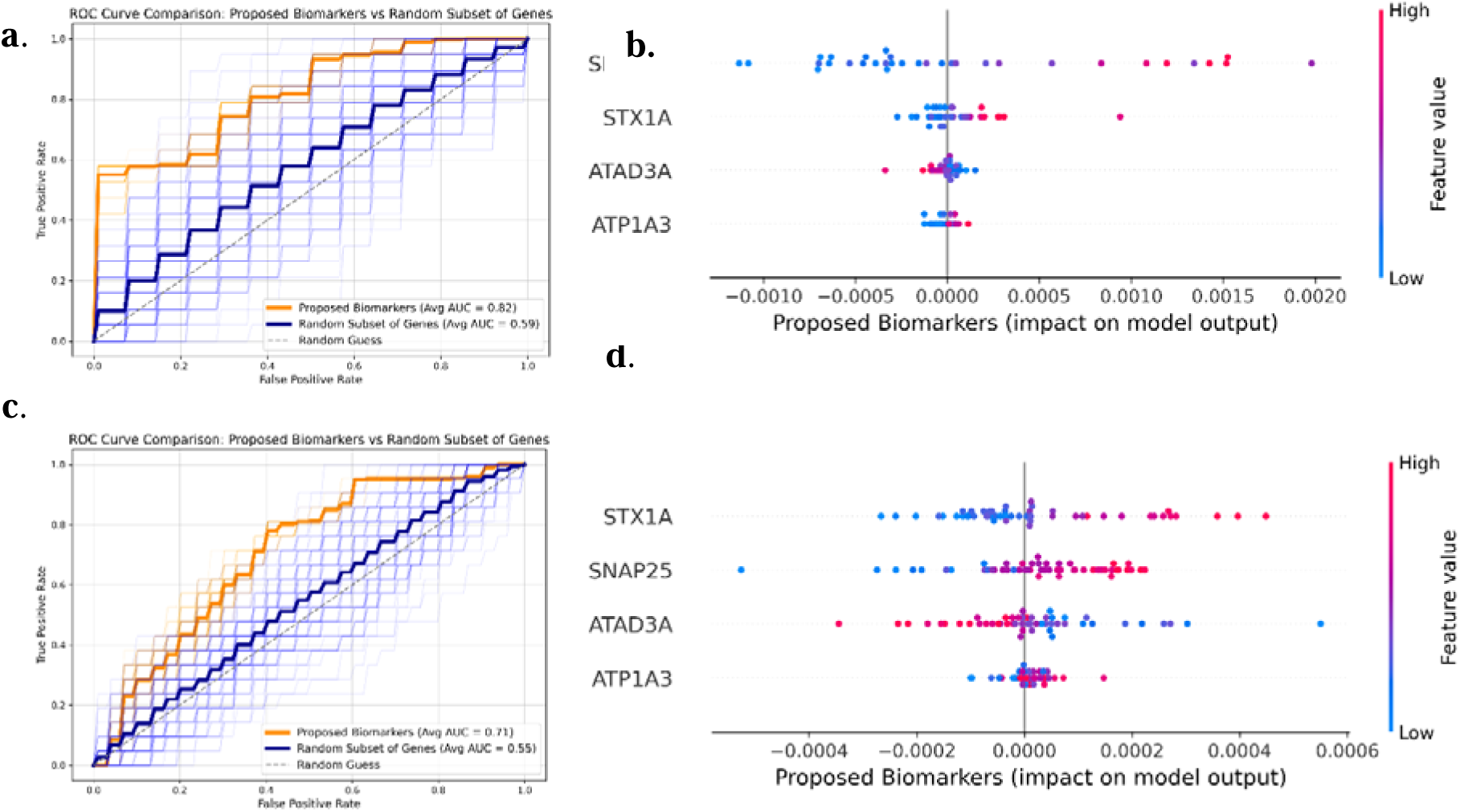
Porosome proteins SNAP-25, Syntaxin-1A and ATP1A3, and the ATAD3A protein essential for mitochondrial fission and bioenergetics, provide superior classification performance in diagnosing AD over other gene products. (a) Receiver Operating Characteristic (ROC) curves for GSE5281, comparing model performance using selected biomarkers (ATP1A3, ATAD3A, SNAP25, STX1A) against randomly selected gene subsets. The model was run 100 times, selecting a random subset of genes in each iteration. The model trained on GSE5281 achieved an AUC of 0.82, significantly outperforming random gene subsets (AUC = 0.59), demonstrating the improved predictive capacity of the identified biomarkers. (b) SHAP-based biomarker importance for GSE5281. SHAP analysis confirms SNAP25 as the top predictor, with other biomarkers contributing to classification. Each gene’s SHAP score is derived from a normalized calculation where the total contribution across all genes sums to 1, providing insight into their relative influence on classification. The color gradient represents feature value distribution, with blue indicating lower expression levels and pink indicating higher expression levels, highlighting the impact of expression variability on model predictions. (c) ROC curves for GSE48350, evaluating classification performance on this dataset. The model was run 100 times, selecting a random subset of genes in each iteration. The targeted biomarkers (ATP1A3, ATAD3A, SNAP25, STX1A) yielded an AUC of 0.71, outperforming randomly selected gene subsets (AUC = 0.55), supporting the validity of these markers. (d) SHAP-based biomarker importance for GSE48350. The color gradient follows the same representation as in (b), indicating the relationship between expression levels and classification outcomes.

**Figure S3.**
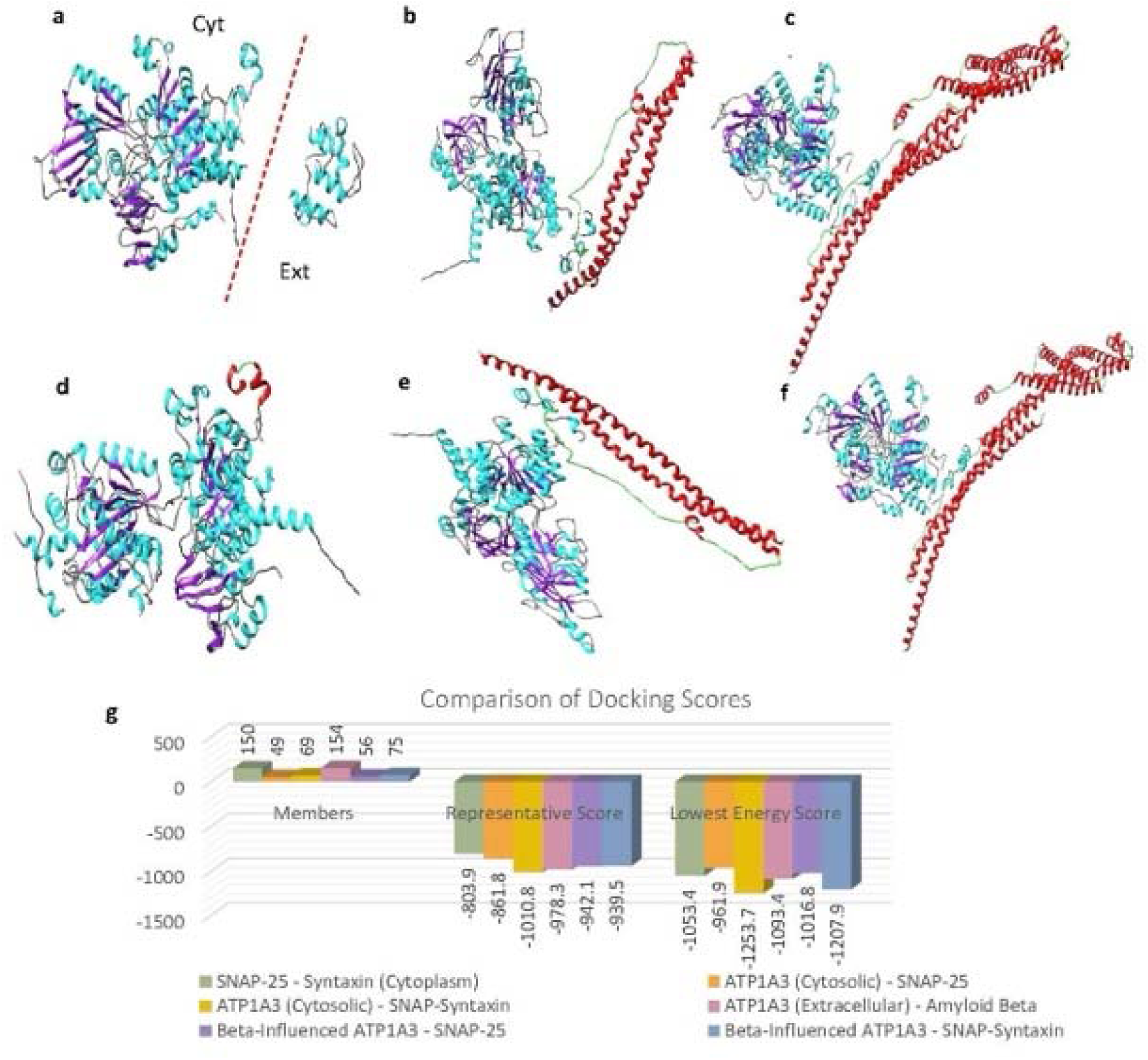
AI reveals altered binding interactions between porosome proteins SNAP-25, Syntaxin-1 and ATP1A3 when amyloid beta 1-42 peptide binding to the extracellular domain of ATP1A3. This binding of the amyloid beta peptide negatively impacts the binding interactions between ATP1A3-SNAP-25-Syntaxin-1A in the cytosolic domain within the neuronal porosome complex. **(a)** Structure of ATP1A3, with cytosolic (Cyt.) and extracellular (Ext.) regions separated by a dashed line to indicate their distinct functional compartments. The extracellular region of ATP1A3 is exposed to the extracellular environment, where it interacts with amyloid beta, while the cytosolic region is involved in intracellular interactions with porosome proteins SNAP-25 and Syntaxin-1. **(b)** The cytosolic domain of ATP1A3 (cyan and purple) favorably binds SNAP-25 (red and green). **(c)** Interaction between the cytosolic domain of ATP1A3 (cyan and purple) and the SNAP-25+Syntaxin-1A complex (red and green), also favorably interact within the porosome machinery. **(d)** Binding of amyloid beta (red and green) to the extracellular region of ATP1A3 (cyan and purple). Amyloid beta binds the extracellular region due to its location outside the cell, where it has been implicated in reducing the ATPase activity of ATP1A3 and neurodegeneration. The image includes the entire ATP1A3 protein, with the cytosolic region added for structural context. **(e)** Binding of amyloid beta to the cytosolic domain of ATP1A3 (cyan and purple) and SNAP-25 (red and green), illustrating potential conformational changes in ATP1A3 induced by amyloid beta binding to its extracellular region. **(f) The b**inding of amyloid beta to the cytosolic domain of ATP1A3 (cyan and purple) alters its binding with the SNAP-25+Syntaxin-1A complex (red and green) within the porosome**. (g)** Comparison of docking scores for various interactions involving ATP1A3, SNAP-25, Syntaxin-1A, and amyloid beta (1-42) peptide. Results show that binding of amyloid beta (1-42) peptide to the extracellular domain of ATP1A3, results in tighter binding (lower energy scores) with SNAP-25 and reduced binding (higher energy scores) with the SNAP-25-Stntaxin-1 complex. This suggests that amyloid beta binding to the extracellular region of ATP1A3 may induce structural changes that enhance intracellular interactions with SNAP-25. These results suggest that while amyloid-beta binding enhances ATP1A3-SNAP-25 interactions, it negatively impacts the formation or stability of the larger ATP1A3-SNAP-25-Syntaxin-1A complex within the neuronal porosome, resulting in impaired secretion of neurotransmitters.

**Figure S4.**
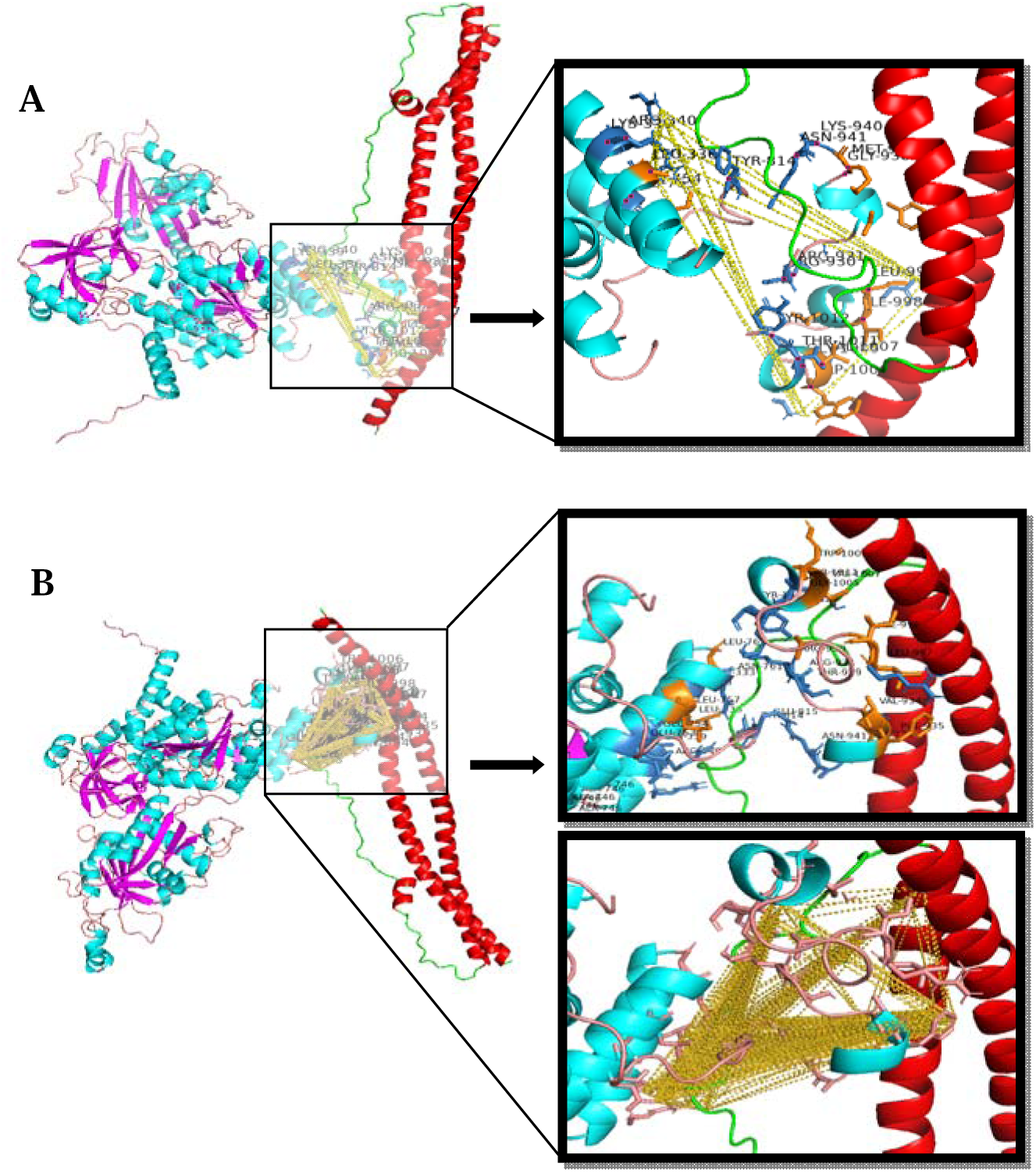
High resolution images of the detailed molecular interaction between ATP1A3 and SNAP25, with and without Amyloid-β binding. This figure illustrates the structural integration of ATP1A3 and SNAP25, highlighting residues within 5Å of the ligand. (A) ATP1A3-SNAP25 Interaction: The interaction without Amyloid-β influence involves 19 key residues (*LEU 336, LYS 339, ARG 340, GLU 753, GLU 754, TYR 814, ARG 930, ARG 931, GLY 938, MET 939, LYS 940, ASN 941, LEU 997, ILE 998, GLY 1005, TRP 1006, VAL 1007, THR 1011, TYR 1012*). (B) ATP1A3-SNAP25 Interaction in the presence of Amyloid-β: With Amyloid-β influence, the number of interacting residues increases to 28, including additional residues such as *CYS 333, LEU 757, ILE 758, ASN 761, LEU 762, GLU 815, THR 929, VAL 934, PHE 935, TYR 1013*. A notable difference in (B) Amyloid-β-influenced ATP1A3-SNAP25 binding is the increase in hydrogen bonds (yellow dashed lines), which were identified based on a 1.8 Å distance threshold. This suggests stronger and potentially altered interactions in the presence of the Amyloid-β peptide. Hydrophilic (skyblue) and hydrophobic (orange) residues are also highlighted, to provide a clearer view, a zoomed-in version of the key residues and hydrogen bonding interactions.

**Figure S5.**
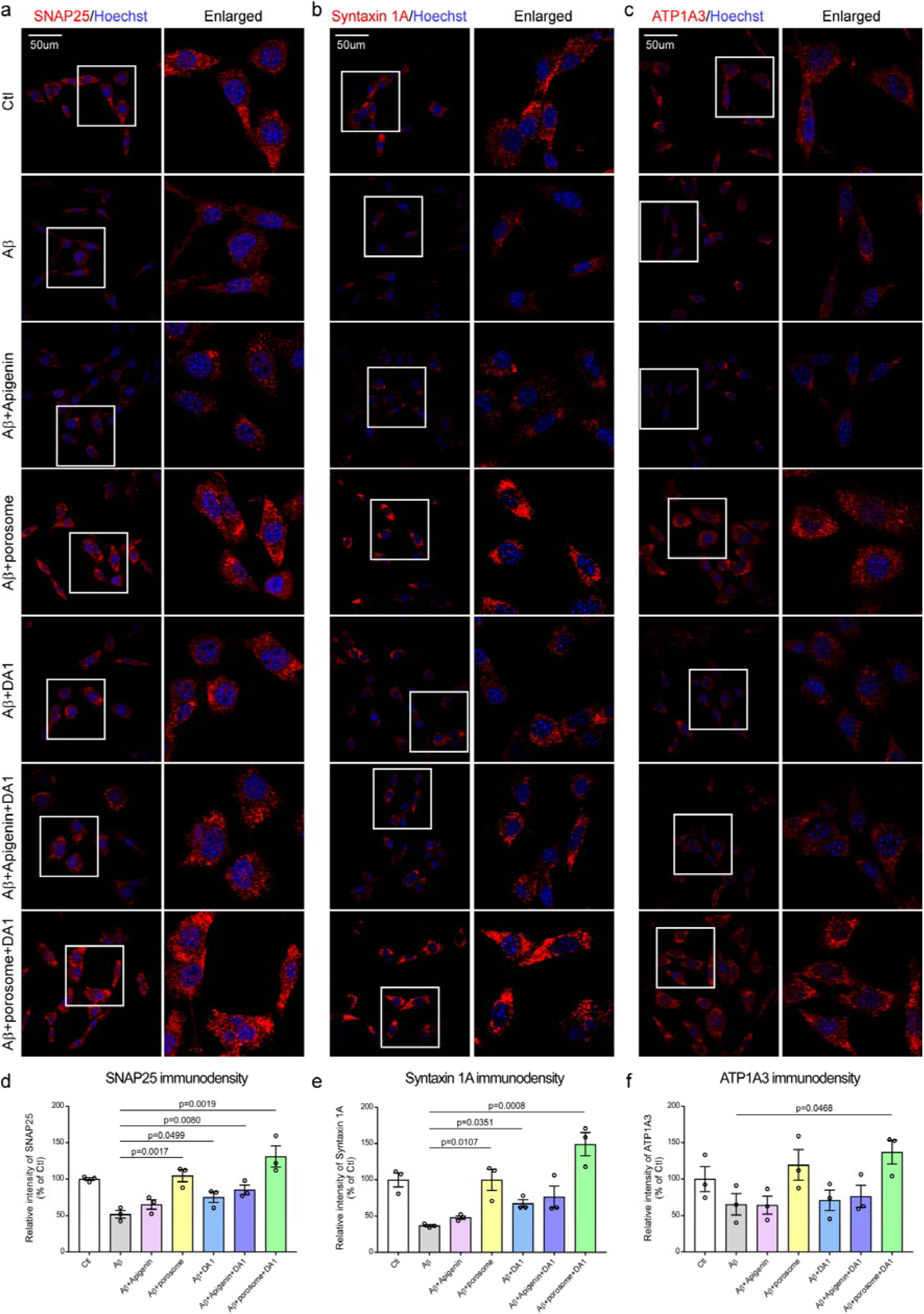
Immunocytochemistry demonstrating depletion of porosome proteins SNAP-25, Syntaxin-1A, and Na^+^/K^+^ Transporting ATPase alpha 3 (ATP1A3) in Alzheimer’s neurons (Aβ treated), which is fully restored to normal levels following porosome-reconstitution therapy, and partially by Apigenin or the DA1 peptide. The combined exposure to both the porosome and DA1 peptide, further enhances the expression of all three porosome proteins: SNAP-25, Syntaxin-1A, and ATP1A3. Note that Apigenin and DA1 combination has a significant effect on enhancing expression of all three porosome proteins. **(a-c)** Mouse hippocampal HT-22 neuronal cells were pretreated with porosome (0.7ug per well in a 24-well plate) overnight. Cells were then pre-incubated with DA1 peptide (1 μM) and Apigenin (5μM) for 1 hour, followed by treatment with oligomeric Aβ_1–42_ peptides (5 μM) for 24 hours. Cells were fixed 1 hour after the second DA1 peptide treatment. Cells were stained with antibodies targeting SNAP25, Syntaxin 1A, and ATP1A3. The immunolocalization of SNAP-25 **(a),** Syntaxin-1A **(b)**, and ATP1A3 **(c)** in untreated control neurons (Ctl), Aβ (Alzheimer’s) neurons (Aβ), Aβ neurons treated with Apigenin (Aβ+Apigenin), Aβ neurons reconstituted with neuronal porosomes (Aβ+porosome), Aβ neurons treated with the DA1 peptide (Aβ+DA1), Aβ neurons treated with Apigenin and the DA1 peptide (Aβ+Apigenin+DA1), and Aβ neurons treated with porosome and the DA1 peptide (Aβ+ porosome+DA1), are shown. **(d-f)** The immunodensities of SNAP-25 **(d)**, Syntaxin-1A **(e)**, and ATP1A3 **(f)** in different treatment groups were quantified and shown in these bar graphs. At least 130 cells per group were analyzed, and the data were from 3 independent experiments. Data are presented as mean ± SEM. Statistical significance was determined by one-way ANOVA with Tukey’s post hoc test.

**Figure S6.**
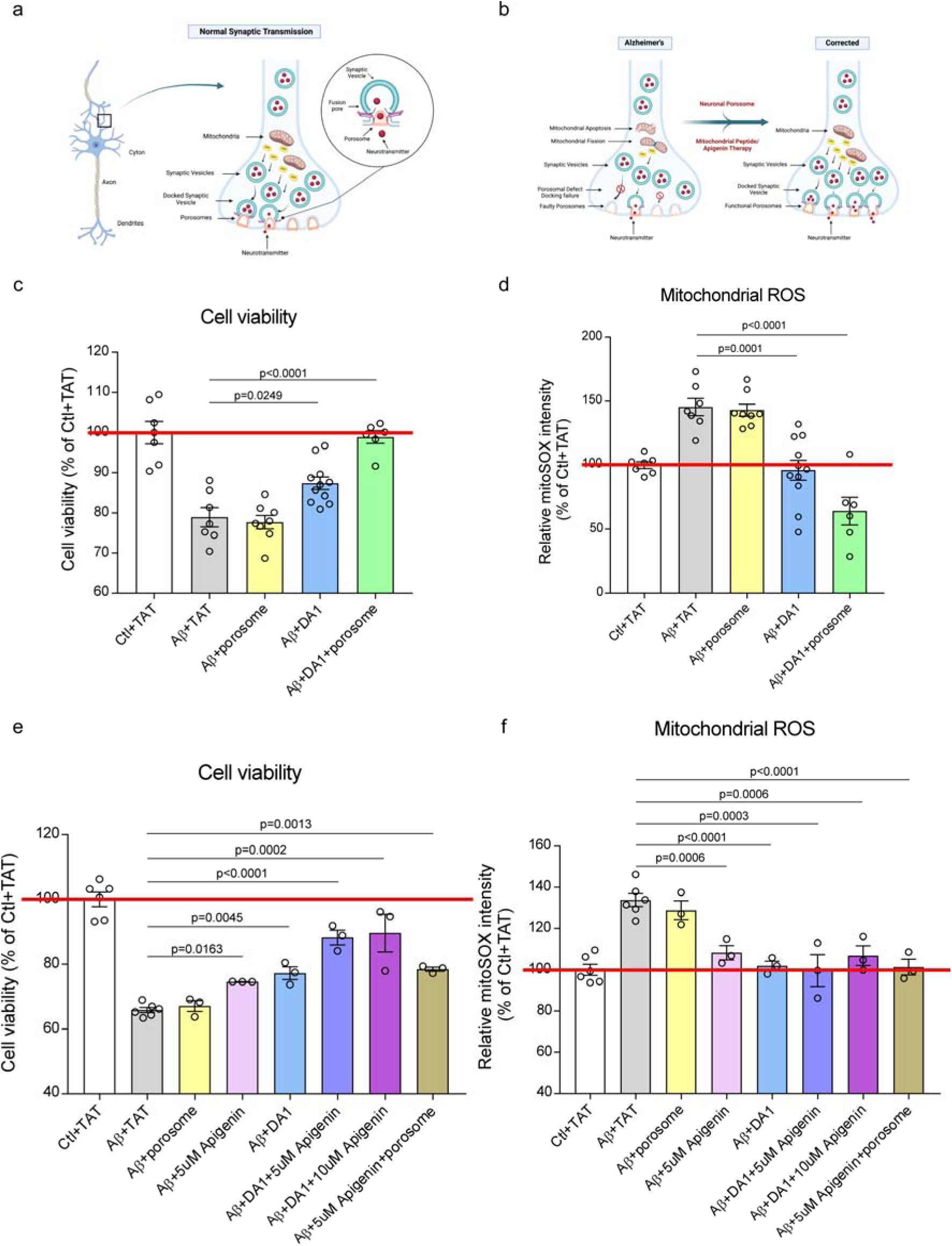
The combinatorial therapy of porosome reconstitution, the secretory corrector, and the DA1 peptide inhibitor of mitochondrial fission, the metabolic corrector, restores cell viability and reduces oxidative stress in mitochondria to normal levels in AD neurons. Similarly, combination therapy of the small flavonoid molecule Apigenin and the mitochondrial fission inhibiting linear DA1 peptide results in increase cell viability and reduction of mitochondrial oxidative stress to near normal levels in AD neurons. **(a)** Schematic illustration of a synapse at the nerve ending in healthy neurons, demonstrating normal mitochondrial function, energy (ATP) generation, and porosome-mediated neurosecretion. **(b)** Schematic illustration of a synapse at the nerve ending in AD neurons, demonstrating abnormal mitochondrial function resulting in mitochondrial fission, loss in ATP generation, and altered porosome-mediated neurosecretion. All of these defects in AD are seen to be overcome using porosome reconstitution and the DA1 peptide inhibitor of mitochondrial fission therapy. **(c)** Cell viability is significantly enhanced in Alzheimer’s neurons (Aβ+TAT) by porosome reconstitution in the presence of the DA1 peptide. Mouse hippocampal HT-22 neuronal cells were pretreated with porosome (175ng per well in a 96-well plate) overnight, followed by treatment with Aβ (oligomeric Aβ_1–42_ peptides) (5μM) together with DA1 (1μM) for 40h. Cell viability was measured by MTT assay after 16h of serum starvation. Note that the cell viability in Alzheimer’s neurons following porosome reconstitution in the presence of the DA1 peptide, demonstrate recovery to healthy neuron levels. Data represent the mean ± SEM from at least six independent experiments. Statistical significance was determined by one-way ANOVA with Tukey’s post hoc test. **(d)** Porosome reconstitution significantly reduces mitochondrial reactive oxygen species (ROS) production in Alzheimer’s neurons (Aβ+TAT) in the presence of DA1 peptide. HT-22 cells were pretreated with porosome (175 ng/well in a 96-well plate) overnight. Cells were then pre-incubated with DA1 (1 μM) for 1 hour, followed by treatment with oligomeric Aβ_1–42_ peptides (10 μM) for 12 hours. Mitochondrial ROS production was assessed 1 hour after the second peptide treatment using MitoSOX staining. MitoSOX intensity in Alzheimer’s neurons following porosome reconstitution in the presence of DA1 peptide demonstrates recovery to much reduced levels than found in healthy neurons. **(e)** Viability of HT-22 cells exposed to 5μM toxic oligomeric Aβ_1–42_ peptides mimicking Alzheimer’s (Aβ+TAT), is significantly enhanced when treated with 1 μM DA1 peptide and 5μM or 10μM Apigenin. Cells were treated with DA1 (1μM) + Aβ (5μM) + Apigenin (5uM or 10μM). After 24h, cells were washed using serum-free media, and again treated with DA1 (1μM) + Aβ (5μM) + Apigenin (5uM or 10μM) in serum-free medium. Cell viability was measured by MTT assay at 40h (16h serum starved). Data represent the mean ± SEM from at least three independent experiments. (**f)** Mitochondrial ROS production in Alzheimer’s neurons (Aβ+TAT) is significantly reduced in the presence of the DA1 peptide. However, not much change is observed in the presence of either 5μM or 10μM of Apigenin. HT-22 cells treated with DA1 (1μM) + Apigenin (5μM or 10μM) for 1h, and then exposed to Aβ (10μM) for 12h. Cells were treated again with DA1 (1uM) + Apigenin (5μM or 10μM) 1h before mitoSOX staining. Data represent the mean ± SEM from at least three independent experiments. Statistical significance was determined by one-way ANOVA with Tukey’s post hoc test.

**Figure S7.**
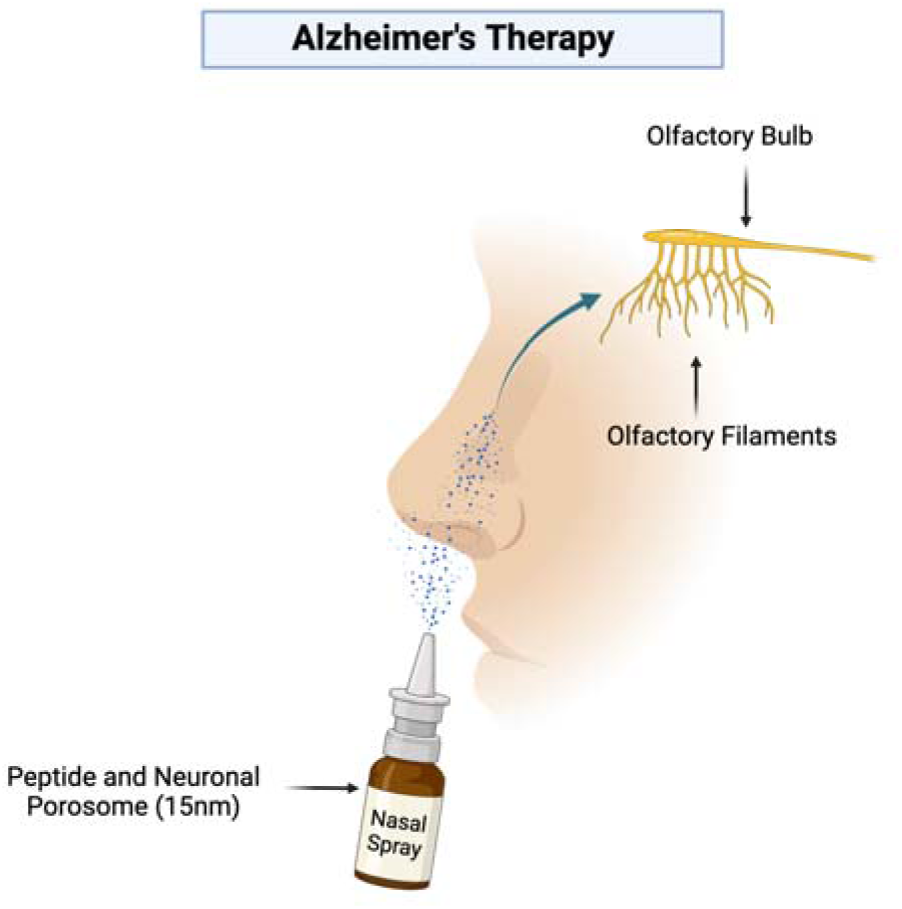
Combination therapy of neuronal porosome or the small flavonoid molecule Apigenin, in combination with the mitochondrial fission inhibiting DA1 peptide for AD therapy. Note that in >90% of Alzheimer’s patients olfactory dysfunction precedes cognitive decline and since the olfactory bulb is connected via nerves to brain regions involved in learning, memory and emotion, this suggests that a combinatorial DA1 peptide therapy either with Apigenin or porosome (a 15 nm biologic), delivered via the nasal route to reach the olfactory bulb, would be most effective in AD therapy, especially at the early stages of the disease.

## References

1. Madeira, C., et al., Elevated Glutamate and Glutamine Levels in the Cerebrospinal Fluid of Patients with Probable Alzheimer’s Disease and Depression. Front Psychiatry, 2018. 9: p. 561.

2. Peña-Bautista, C., et al., Plasma alterations in cholinergic and serotonergic systems in early Alzheimer Disease: Diagnosis utility. Clin Chim Acta, 2020. 500: p. 233–240.

3. Snowden, S.G., et al., Neurotransmitter Imbalance in the Brain and Alzheimer’s Disease Pathology. J Alzheimers Dis, 2019. 72(1): p. 35–43.

4. Swerdlow, R.H., Mitochondria and Mitochondrial Cascades in Alzheimer’s Disease. J Alzheimers Dis, 2018. 62(3): p. 1403–1416.

5. Yu, Q., et al., Antioxidants Rescue Mitochondrial Transport in Differentiated Alzheimer’s Disease Trans-Mitochondrial Cybrid Cells. J Alzheimers Dis, 2016. 54(2): p. 679–90.

6. Pickett, E.K., et al., Region-specific depletion of synaptic mitochondria in the brains of patients with Alzheimer’s disease. Acta Neuropathol, 2018. 136(5): p. 747–757.

7. Jumper, J., et al., Highly accurate protein structure prediction with AlphaFold. Nature, 2021. 596(7873): p. 583–589.

8. Jena, Bhanu P. Secretion machinery at the cell plasma membrane. Current opinion in structural biology vol. 17,4:437–43. doi: 10.1016/j.sbi.2007.07.002 (2007).

9. Schekman, Randy. The Secrets of Secretion: Protein Transport in Cells. Frontiers for Young Minds. 11. 10.3389/frym.2023.1063926. (2023).

10. Lee, J.S., et al., Neuronal porosome proteome: Molecular dynamics and architecture. J Proteomics, 2012. 75(13): p. 3952–62.

11. Zhao, Y., et al., ATAD3A oligomerization promotes neuropathology and cognitive deficits in Alzheimer’s disease models. Nat Commun, 2022. 13(1): p. 1121.

12. Zhao, Y., et al., ATAD3A oligomerization causes neurodegeneration by coupling mitochondrial fragmentation and bioenergetics defects. Nat Commun, 2019. 10(1): p. 1371.

13. World Alzheimer Report 2016, Alzheimer’s Disease International. Lincolnshire, IL. (2016).

14. Reddy, P.H., et al., Abnormal mitochondrial dynamics and synaptic degeneration as early events in Alzheimer’s disease: implications to mitochondria-targeted antioxidant therapeutics. Biochim Biophys Acta, 2012. 1822(5): p. 639–49.

15. Reddy, P.H. and S. McWeeney, Mapping cellular transcriptosomes in autopsied Alzheimer’s disease subjects and relevant animal models. Neurobiol Aging, 2006. 27(8): p. 1060–77.

16. Area-Gomez, E., et al., A key role for MAM in mediating mitochondrial dysfunction in Alzheimer disease. Cell Death Dis, 2018. 9(3): p. 335.

17. Mutisya, E.M., A.C. Bowling, and M.F. Beal, Cortical cytochrome oxidase activity is reduced in Alzheimer’s disease. J Neurochem, 1994. 63(6): p. 2179–84.

18. Adlimoghaddam, A., et al., Regional hypometabolism in the 3xTg mouse model of Alzheimer’s disease. Neurobiol Dis, 2019. 127: p. 264–277.

19. Cao, L., et al., *A*β *alters the connectivity of olfactory neurons in the absence of amyloid plaques in vivo*. Nat Commun, 2012. 3: p. 1009.

20. Wang, Q., et al., SNAP25 is a potential target for early stage Alzheimer’s disease and Parkinson’s disease. Eur J Med Res, 2023. 28(1): p. 570.

21. Reinikainen, K.J., A. Pitkänen, and P.J. Riekkinen, 2’,3’-cyclic nucleotide-3’-phosphodiesterase activity as an index of myelin in the post-mortem brains of patients with Alzheimer’s disease. Neurosci Lett, 1989. 106(1-2): p. 229–32.

22. Sinclair, L.I., H.M. Tayler, and S. Love, Synaptic protein levels altered in vascular dementia. Neuropathol Appl Neurobiol, 2015. 41(4): p. 533–43.

23. Mukaetova-Ladinska, E.B., et al., Lewy body variant of Alzheimer’s disease: selective neocortical loss of t-SNARE proteins and loss of MAP2 and alpha-synuclein in medial temporal lobe. ScientificWorldJournal, 2009. 9: p. 1463–75.

24. Greber, S., et al., Decreased levels of synaptosomal associated protein 25 in the brain of patients with Down syndrome and Alzheimer’s disease. Electrophoresis, 1999. 20(4-5): p. 928–34.

25. Corradini, I., et al., Epileptiform activity and cognitive deficits in SNAP-25(+/-) mice are normalized by antiepileptic drugs. Cereb Cortex, 2014. 24(2): p. 364–76.

26. Won Jin, C., et al., Porosome reconstitution therapy: A biologic rescue from cystic fibrosis. bioRxiv, 2024: p. 2024.09.11.612494.

27. Murphy, M.P., How mitochondria produce reactive oxygen species. Biochem J, 2009. 417(1): p. 1–13.

28. Lin, M.T. and M.F. Beal, Mitochondrial dysfunction and oxidative stress in neurodegenerative diseases. Nature, 2006. 443(7113): p. 787–95.

29. Bors, W., C. Michel, and M. Saran, Flavonoid antioxidants: rate constants for reactions with oxygen radicals. Methods Enzymol, 1994. 234: p. 420–9.

30. Pannala, A.S., et al., Inhibition of peroxynitrite-mediated tyrosine nitration by catechin polyphenols. Biochem Biophys Res Commun, 1997. 232(1): p. 164–8.

31. Heijnen, C.G., et al., Flavonoids as peroxynitrite scavengers: the role of the hydroxyl groups. Toxicol In Vitro, 2001. 15(1): p. 3–6.

32. Gilquin, B., et al., The AAA+ ATPase ATAD3A controls mitochondrial dynamics at the interface of the inner and outer membranes. Mol Cell Biol, 2010. 30(8): p. 1984–96.

33. Issop, L., et al., Mitochondria-associated membrane formation in hormone-stimulated Leydig cell steroidogenesis: role of ATAD3. Endocrinology, 2015. 156(1): p. 334–45.

34. Baudier, J., ATAD3 proteins: brokers of a mitochondria-endoplasmic reticulum connection in mammalian cells. Biol Rev Camb Philos Soc, 2018. 93(2): p. 827–844.

35. Chiang, S.F., et al., An alternative import pathway of AIF to the mitochondria. Int J Mol Med, 2012. 29(3): p. 365–72.

36. Goller, T., et al., Atad3 function is essential for early post-implantation development in the mouse. PLoS One, 2013. 8(1): p. e54799.

37. Lee, H. and D.W. Kim, Deletion of ATAD3A inhibits osteogenesis by impairing mitochondria structure and function in pre-osteoblast. Dev Dyn, 2022. 251(12): p. 1982–2000.

38. Lang, L., et al., ATAD3A mediates activation of RAS-independent mitochondrial ERK1/2 signaling, favoring head and neck cancer development. J Exp Clin Cancer Res, 2022. 41(1): p. 43.

39. Harel, T., et al., Recurrent De Novo and Biallelic Variation of ATAD3A, Encoding a Mitochondrial Membrane Protein, Results in Distinct Neurological Syndromes. Am J Hum Genet, 2016. 99(4): p. 831–845.

40. Cooper, H.M., et al., ATPase-deficient mitochondrial inner membrane protein ATAD3A disturbs mitochondrial dynamics in dominant hereditary spastic paraplegia. Hum Mol Genet, 2017. 26(8): p. 1432–1443.

41. Hoerder-Suabedissen, A., et al., Cell-Specific Loss of SNAP25 from Cortical Projection Neurons Allows Normal Development but Causes Subsequent Neurodegeneration. Cereb Cortex, 2019. 29(5): p. 2148–2159.

42. Kivisäkk, P., et al., Increased levels of the synaptic proteins PSD-95, SNAP-25, and neurogranin in the cerebrospinal fluid of patients with Alzheimer’s disease. Alzheimers Res Ther, 2022. 14(1): p. 58.

43. Cummings, J., et al., Alzheimer’s disease drug development pipeline: 2023. Alzheimers Dement (N Y), 2023. 9(2): p. e12385.

44. Vyhnalek, M., et al., Olfactory identification in amnestic and non-amnestic mild cognitive impairment and its neuropsychological correlates. J Neurol Sci, 2015. 349(1-2): p. 179–84.

45. Halbgebauer, S., et al., CSF levels of SNAP-25 are increased early in Creutzfeldt-Jakob and Alzheimer’s disease. J Neurol Neurosurg Psychiatry, 2022.

46. Woodward, M.R., et al., Odorant Item Specific Olfactory Identification Deficit May Differentiate Alzheimer Disease From Aging. Am J Geriatr Psychiatry, 2018. 26(8): p. 835–846.

47. Xiong, H., et al., Cholesterol retention in Alzheimer’s brain is responsible for high beta- and gamma-secretase activities and Abeta production. Neurobiol Dis, 2008. 29(3): p. 422–37.

48. Schrödinger, LLC. PyMOL: The PyMOL Molecular Graphics System, Version 2.0. (2015).

49. Kozakov, D., et al., The ClusPro web server for protein-protein docking. Nat Protoc, 2017. 12(2): p. 255–278.

50. Won Jin, C., et al., Reprogramming the neuronal secretory and metabolic machinery using correctors for Alzheimer therapy. bioRxiv, 2024: p. 2024.09.11.622805.

